# *In vitro* screening of a FDA approved chemical library reveals potential inhibitors of SARS-CoV-2 replication

**DOI:** 10.1101/2020.04.03.023846

**Authors:** Franck Touret, Magali Gilles, Karine Barral, Antoine Nougairède, Etienne Decroly, Xavier de Lamballerie, Bruno Coutard

## Abstract

A novel coronavirus, named SARS-CoV-2, emerged in 2019 from Hubei region in China and rapidly spread worldwide. As no approved therapeutics exists to treat Covid-19, the disease associated to SARS-Cov-2, there is an urgent need to propose molecules that could quickly enter into clinics. Repurposing of approved drugs is a strategy that can bypass the time consuming stages of drug development. In this study, we screened the Prestwick Chemical Library® composed of 1,520 approved drugs in an infected cell-based assay. 90 compounds were identified. The robustness of the screen was assessed by the identification of drugs, such as Chloroquine derivatives and protease inhibitors, already in clinical trials. The hits were sorted according to their chemical composition and their known therapeutic effect, then EC50 and CC50 were determined for a subset of compounds. Several drugs, such as Azithromycine, Opipramol, Quinidine or Omeprazol present antiviral potency with 2<EC50<20µM. By providing new information on molecules inhibiting SARS-CoV-2 replication *in vitro*, this study could contribute to the short-term repurposing of drugs against Covid-19.

## Introduction

Human coronaviruses (HCoVs) are enveloped positive-stranded RNA viruses belonging to the *Nidovirales* order that are mostly involved in gastrointestinal and respiratory tract infections. Among them, severe acute respiratory syndrome (SARS) and Middle East respiratory syndrome (MERS) CoVs that emerged in 2002 and 2013 respectively, have been associated with severe human illnesses, such as severe acute respiratory distress syndromes (de Wit et al., 2016). In December 2019, a new coronavirus (SARS-CoV-2) has emerged in the city of Wuhan and quickly spread around the world. SARS-CoV-2 causes in human a viral infection, named COVID-19, which is associated in some patients with severe respiratory diseases and significant mortality rates, in particular in elderly populations (Li et al., 2020). While an unknown fraction (most probably a majority) of infected people remain pauci- or asymptomatic, some require hospitalization, sometimes in intensive care units, which may jeopardise health systems during peak pandemic periods. In such a context, vaccines would represent great tools to prevent or limit virus spread. However, vaccine development is a long process and COVID-19 vaccines will most probably not be concretely available for mass usage, at least during the first wave of the disease. Accordingly, the availability of efficient antiviral drugs would be of utmost interest for the treatment of infected patients and possibly for preventive or preemptive use. Regrettably, the current and unprecedented outbreak of SARS-CoV-2 occurs in an unprepared world, with no firmly established identification of active molecules against beta-coronaviruses (Callaway et al., 2020). There is thus an urgent necessity to provide *hic et nunc* therapeutic solutions to limit viral infection. As the timeframe for a conventional drug development is unrelated to the immediate needs, repurposing of drugs originally developed for other viral infections or therapeutic uses is likely the fastest way to enter clinics. This fast track drug development and validation lead to the initiation of numerous clinical trials for the treatment of Covid19 (Li and De Clercq, 2020) but there is still a need to expand the number of possible drug candidates to treat COVID19 and/or evaluate possible drug combinations to potentiate the antiviral effects (Cheng et al., 2019). Whereas the number of clinical trials cannot be extensively multiplied, libraries of “old” drugs can be screened *in vitro* in medium to high throughput assays. Proof of concepts for drug repositioning against SARS-CoV, MERS-CoV or other viruses in *in vitro* assay already showed the relevance of this strategy (de Wilde et al., 2014; Dyall et al., 2014; Mercorelli et al., 2018). In this study, we screened the 1,520 approved and off-patent drugs of the Prestwick Chemical Library^®^ in a SARS-CoV-2 infection cell-based assay. The *in vitro* screening identified 90 drugs showing inhibition effect on the viral replication at 10 μM. Hits, selected from different classes of compounds, were then confirmed by EC50 and CC50 determination.

## Material and methods

### Chemical library

The Prestwick Chemical Library® (hereafter named PCL) is a library of 1,520 off-patent small molecules, mostly approved drugs (FDA, EMA and other agencies). The compounds are provided at a concentration of 10mM in 100% DMSO.

### Cell line

VeroE6 (ATCC CRL-1586) cells were grown in minimal essential medium (Life Technologies) with 7.5% heat-inactivated fetal calf serum (FCS), at 37°C with 5% CO^2^ with 1% penicillin/streptomycin (PS, 5000U.mL^−1^ and 5000µg.mL^−1^ respectively; Life Technologies) and supplemented with 1 % non-essential amino acids (Life Technologies).

### Virus strain

SARS-CoV-2 strain BavPat1 was obtained from Pr Drosten through EVA GLOBAL (https://www.european-virus-archive.com/). To prepare the virus working stock, a 25cm^2^ culture flask of confluent VeroE6 cells growing with MEM medium with 2.5% FBS (Life Technologies) was inoculated at MOI 0.001. Cell supernatant medium was harvested at the peak of infection and supplemented with 25mM HEPES (Sigma) before being stored frozen in small aliquots at −80°C. All experiments were conducted in BSL3 laboratory.

### Antiviral screen

One day prior to infection for the antiviral screening 5×10^4^ VeroE6 cells were seeded in 100µL assay medium (containing 2.5% FCS) in 96 well plates. The next day, a single dilution of each compound of the PCL at 10µM final concentration was added to the cells (25µL/well, in 2.5% FCS-containing medium). Six virus control wells were supplemented with 25µL medium (positive controls hereafter named vc) and eight cell control wells were supplemented with 50µL of medium (negative controls, hereafter named nc). Two internal well controls of viral inhibition were added, corresponding to the addition of 10µM arbidol (Sigma) in the infected cell culture (arbidol controls, hereafter named arb). After 15 min, 25µL of a virus mix diluted in 2.5% FCS-containing medium was added to the wells at MOI 0.002.

Three days after infection, cell supernatant media were discarded and CellTiter-Blue® reagent (Promega) was added following the manufacturer’s instructions. Plates were incubated for 2 hours prior recording fluorescence (560/590nm) with a Tecan Infinite 200Pro machine. From the measured OD_590nm_, the Inhibition Index was calculated as follows: first, cell viability for compounds, vc and arb were calculated: (OD_590nm_ value/mean OD_590nm_ of nc)*100. For vc and arb, mean cell viability were calculated. Then all cell viabilities were normalized by subtracting mean vc. cell viability of the 96 well plates. Finally, Inhibition index was calculated as follows: Inh. Index= normalized cell viability of the compound/normalized cell viability of arb in the same 96 well plate. All compounds with Inhibition index values above 1 were considered as a hit.

### EC50 and CC50 determination

One day prior to infection, 5×104 VeroE6 cells were seeded in 100µL assay medium (containing 2.5% FCS) in 96 well plates. The next day, seven 2-fold serial dilutions of compounds (0.6µM to 40µM, in triplicate) were added to the cells (25µL/well, in assay medium). Four virus control wells were supplemented with 25µL of assay medium. After 15 min, 25µL of a virus mix diluted in medium was added to the wells. The amount of virus working stock used was calibrated prior to the assay, based on a replication kinetics, so that the replication growth is still in the exponential growth phase for the readout as previously described (Delang et al., 2016; Touret et al., 2019). Four cell control wells (*i*.*e*. with no virus) were supplemented with 50µL of assay medium. On each plate a control compound (Remdesivir, BLDpharm) was added in duplicate with seven 2-fold serial dilutions (0.16µM to 20µM, in duplicate). Plates were incubated for 2 days at 37°C prior to quantification of the viral genome by real-time RT-PCR. To do so, 100µL of viral supernatant was collected in S-Block (Qiagen) previously loaded with VXL lysis buffer containing proteinase K and RNA carrier. RNA extraction was performed using the Qiacube HT automat and the Cador Pathogen 96 HT kit following manufacturer instruction. Viral RNA was quantified by real-time RT-qPCR (EXPRESS One-Step Superscript™ qRT-PCR Kit, universal Invitrogen using 3.5µL of RNA and 6.5µL of RT qPCR mix and standard fast cycling parameters, *i*.*e*., 10min at 50°C, 2 min at 95°C, and 40 amplification cycles (95°C for 3 sec followed by 30sec at 60°C). Quantification was provided by four 2 log serial dilutions of an appropriate T7-generated synthetic RNA standard of known quantities (10^2^ to 10^8^ copies). RT-qPCR reactions were performed on QuantStudio 12K Flex Real-Time PCR System (Applied Biosystems) and analyzed using QuantStudio 12K Flex Applied Biosystems software v1.2.3. Primers and probe sequences, which target SARS-CoV-2 N gene, were: Fw: GGCCGCAAATTGCACAAT; Rev : CCAATGCGCGACATTCC; Probe: FAM-CCCCCAGCGCTTCAGCGTTCT-BHQ1. The 50% and 90% effective concentrations (EC50, EC90; compound concentration required to inhibit viral RNA replication by 50% and 90%) were determined using logarithmic interpolation as previously described (Touret et al., 2019). For the evaluation of the CC50 (the concentration that reduces the total cell number by 50%), the same culture conditions were set as for the determination of the EC50, without addition of the virus, and cell viability was measured using CellTiter Blue® (Promega) as previously described for the screening. CC50 was determined using logarithmic interpolation. All data obtained were analyzed using Graph pad prism 7 software (Graph pad software). All graphical representations were also performed on Graph pad prism 7 software.

## Results and discussion

We developed a HTS SARS-CoV-2 replication inhibition assay based on the measurement of the cell viability 3 days after cell infection with a MOI of 0.002. Prior to the screening, we evaluated the antiviral effect of arbidol, a broad spectrum antiviral compounds that blokes the viral entry of many enveloped viruses (Blaising et al., 2014). In our experimental conditions, we demonstrated that 10µM arbidol limit the SARS-CoV-2 infection leading to around 70-90% cell viability, with a EC50 of 10.7µM. This compound was next used as reference compounds in order to calculate the inhibition index (Inh. Index).

We next tested the Prestwick Chemical Library® (PCL®) composed of 1,520 approved drugs at a final concentration of 10µM. The cell viability was determined and we calculated the relative value of inhibition potency compared to arbidol. Among the 1,520 compounds of the PCL®, 90 compounds showed equal or more potent inhibition than arbidol with an Inh. Index ≥ 1 (5.85 % positive hits) (Fig.1), and the mean Inh. Index of the library was 0.28 (Table 1 of the supplemental data). As the threshold for the selection is arbitrary, the raw data for each compound of the PCL® is presented in the supplemental data, in order to allow the scientific community to further analyse the results and possibly rescue molecules of interest.

**Figure 1:**
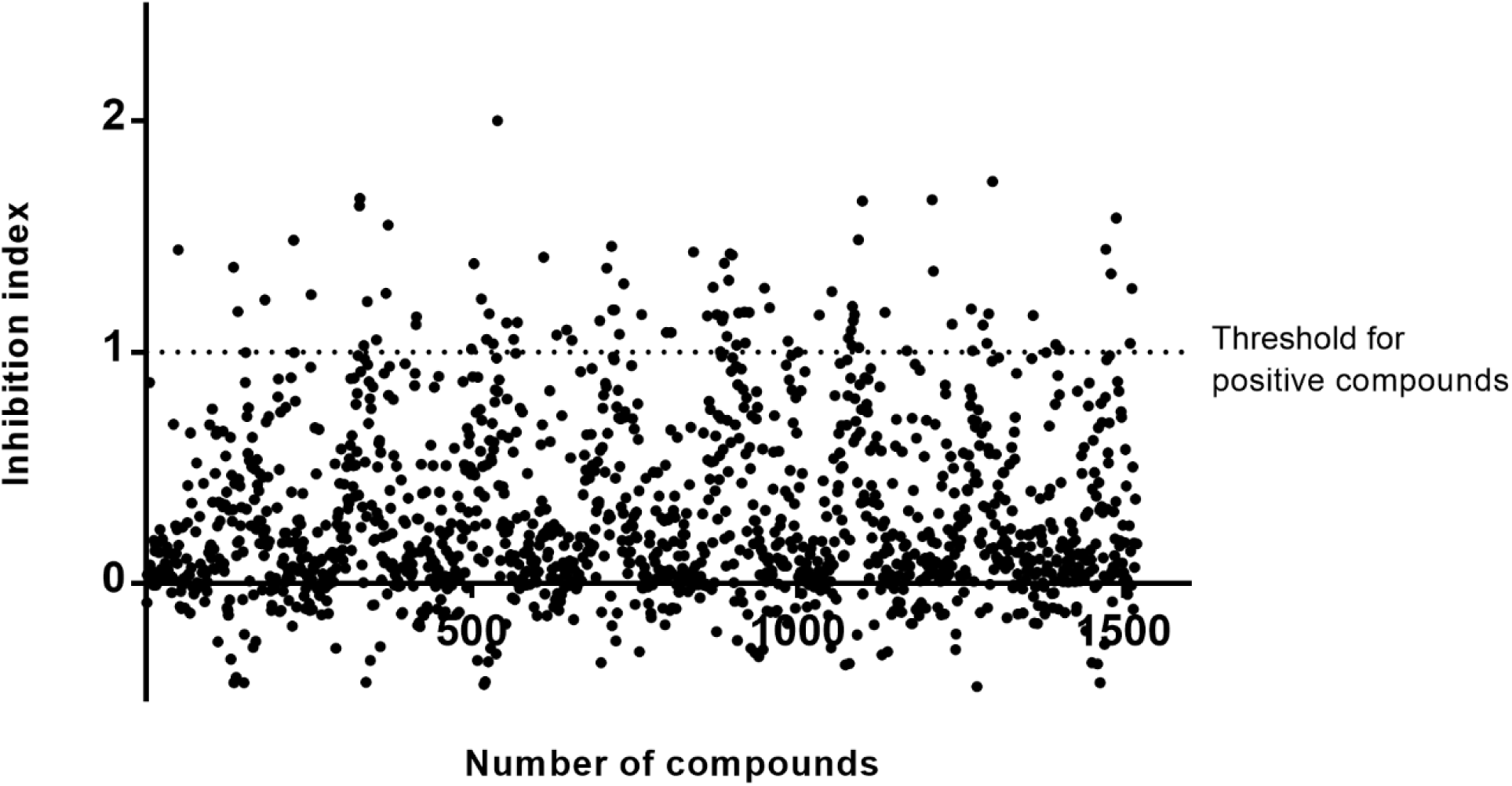
Screening of 1,520 clinically approved compounds from Prestwick Chemical Library® and hits selection. The threshold was based on the *in vitro* antiviral potency of arbidol at 10µM.

**Table 1:** Antiviral activity of selected compounds from the hits and control compounds. EC50: 50% inhibition, EC90: 90% inhibition, CC50:50% cytotoxicity and SI: selectivity index. n.d: not determined. All value are in µM. *From (Haviernik et al., 2018)

|  | EC50 | EC90 | CC50 | SI |
| --- | --- | --- | --- | --- |
| **Azithromycine** | 2,12 | 8,65 | >40 | >19 |
| **Omeprazole** | 17,06 | 38,01 | >40 | >2,35 |
| **Oxprenolol hydrochloride** | 20,22 | >40 | >40 | >2 |
| **Hydroxy-chloroquine** | 4,17 | 25,49 | >40 | >10 |
| **Clemizole hydrochloride** | 23,94 | 38,23 | >40 | >1,7 |
| **Alprostadil** | 5,39 | 62,40 | >40 | >7,4 |
| **Dolutegravir** | 22,04 | 42,81 | >40 | >1,8 |
| **Sulfadoxine** | 35,37 | 45,11 | >40 | >1,13 |
| **Opipramol dihydrochloride** | 5,05 | 5,97 | >40 | >7,9 |
| **Quinidine hydrochloride monohydrate** | 5,11 | >40 | >40 | >7,8 |
| **Arbidol** | 10,7 | 15,2 | >40* | >3,7 |
| **Remdesivir 5 exp** | 1,65±0,79 | 2,52±0,81 | n.d | n.d |

Among the selected hits, we first identified drugs previously demonstrated to inhibit *in vitro* the SAR-CoV2 replication. Accordingly, Chloroquine and Hydroxychloroquine (Liu et al., 2020; Wang et al., 2020; Yao et al., 2020) were shown to limit SARS-CoV-2 replication with a Inh. Index of 1.35 and 1.43 respectively. In addition to chloroquine derivatives, two hits are also currently evaluated in different clinical trials, namely Darunavir and Azythromycine. These observations assessed the robustness of the screening assay. In order to further consolidate the results provided by the assay, EC50 was determined on a set of selected compounds. Whereas the screening relied on the quantification of cytopathic effect (CPE) using CellTiter Blue® providing qualitative information on the viral infection, EC50 determination were based on the quantification of the viral genome by Real-Time RT-PCR (Touret et al., 2019). For this assay Remdesivir was used as a control compound for validation, with an EC50 of 1.6µM (Table 1), a value in agreement with previously published data (Wang et al., 2020).

From the 90 selected hits (Table 2), some were arbitrary removed after visual inspection of their initial therapeutic use or strong side effects (ophthalmic treatment, topical administration, teratogenic effect …). The compounds were organised in 12 groups by structural similarity and/or therapeutic class including ten steroids (mainly sex steroids used for endocrine therapy and corticosteroids with anti-inflammatory properties), two prostaglandins (both PGE1 displaying a variety of pharmacologic actions such as vasodilatator for one and anti-ulcer for the other), two proton pump inhibitors (PPIs used as anti-ulcer agents), two antiretrovirals (HIV anti-protease and anti-integrase), twelve antibacterial drugs (including fluoroquinolone, oxazolidinone, macrolide and beta-lactam derivatives among others), fifteen cardiovascular drugs (antiarrhythmic, vasodilatator and antihypertensive agents including ion channel blockers, an angiotensin converting enzyme inhibitor (ACE), angiotensin II receptor inhibitors, and beta-adrenergic receptor blockers), two opioids plus sixteen non-opioid CNS drugs (that display a variety of pharmacologic actions such as antipsychotic, antidepressant, antiparkinsonian, anti-alzheimer…), six neuromuscular-blocking drugs (including muscle relaxant and local anesthesic agents), three respiratory system drugs (mostly used as a bronchodilator in the treatment of asthma and COPD), three allergy medications (with antihistaminic and antiemetic properties), three antiparasitic drugs (more specifically antimalarial agents), and fourteen unrelated drugs (used in ophthalmology, dermatology and other isolated pathologies).

**Table 2:**
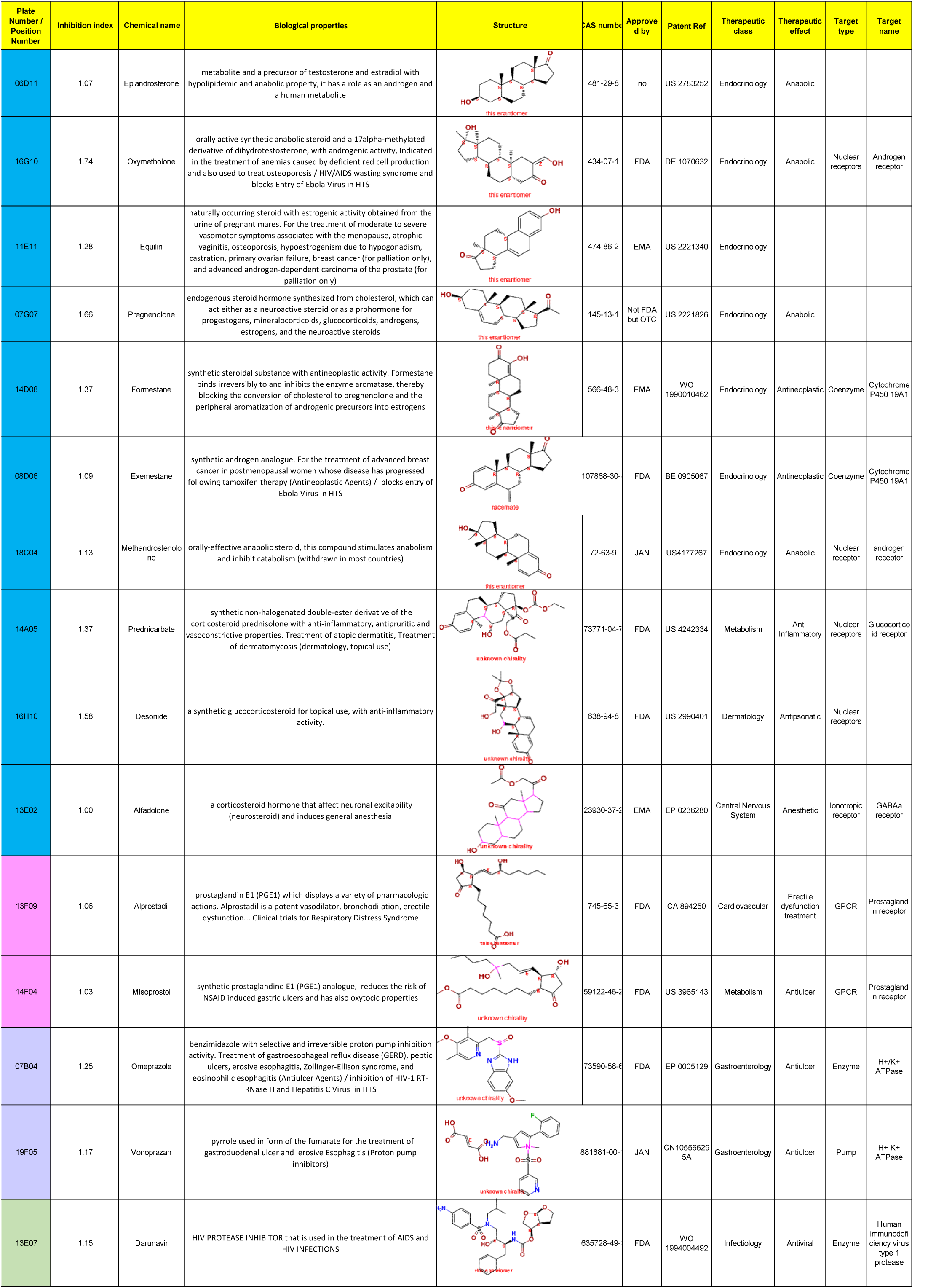
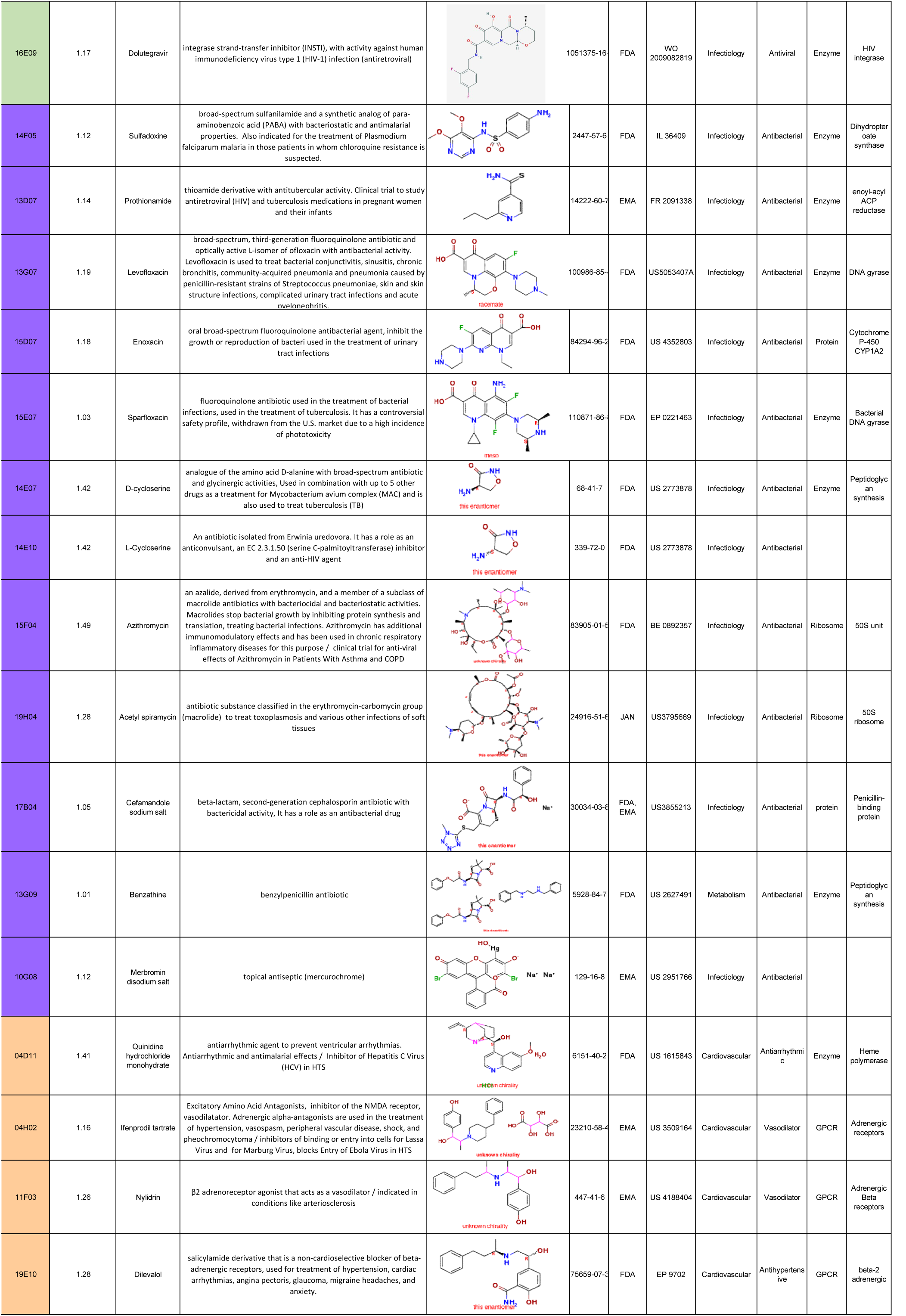
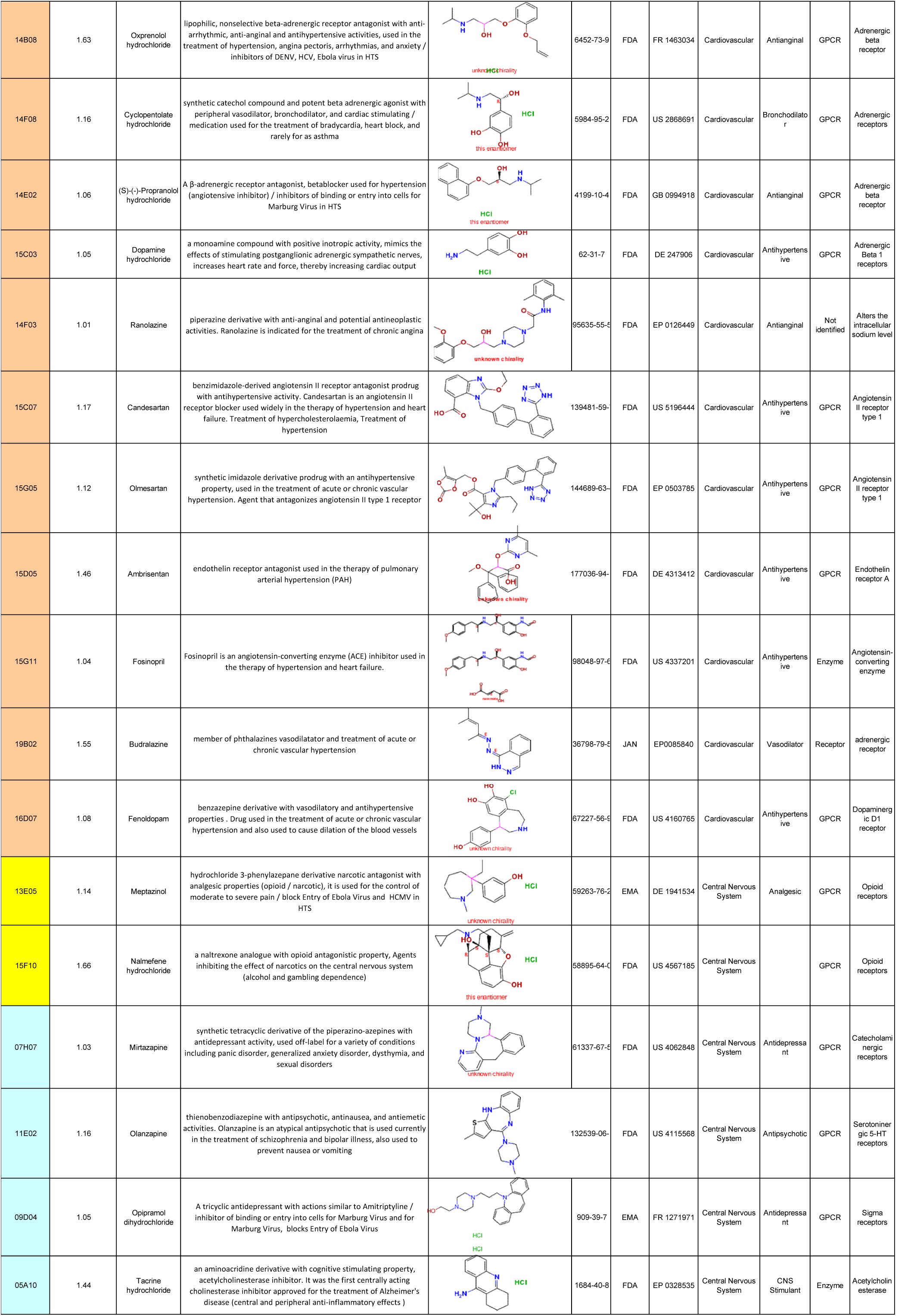
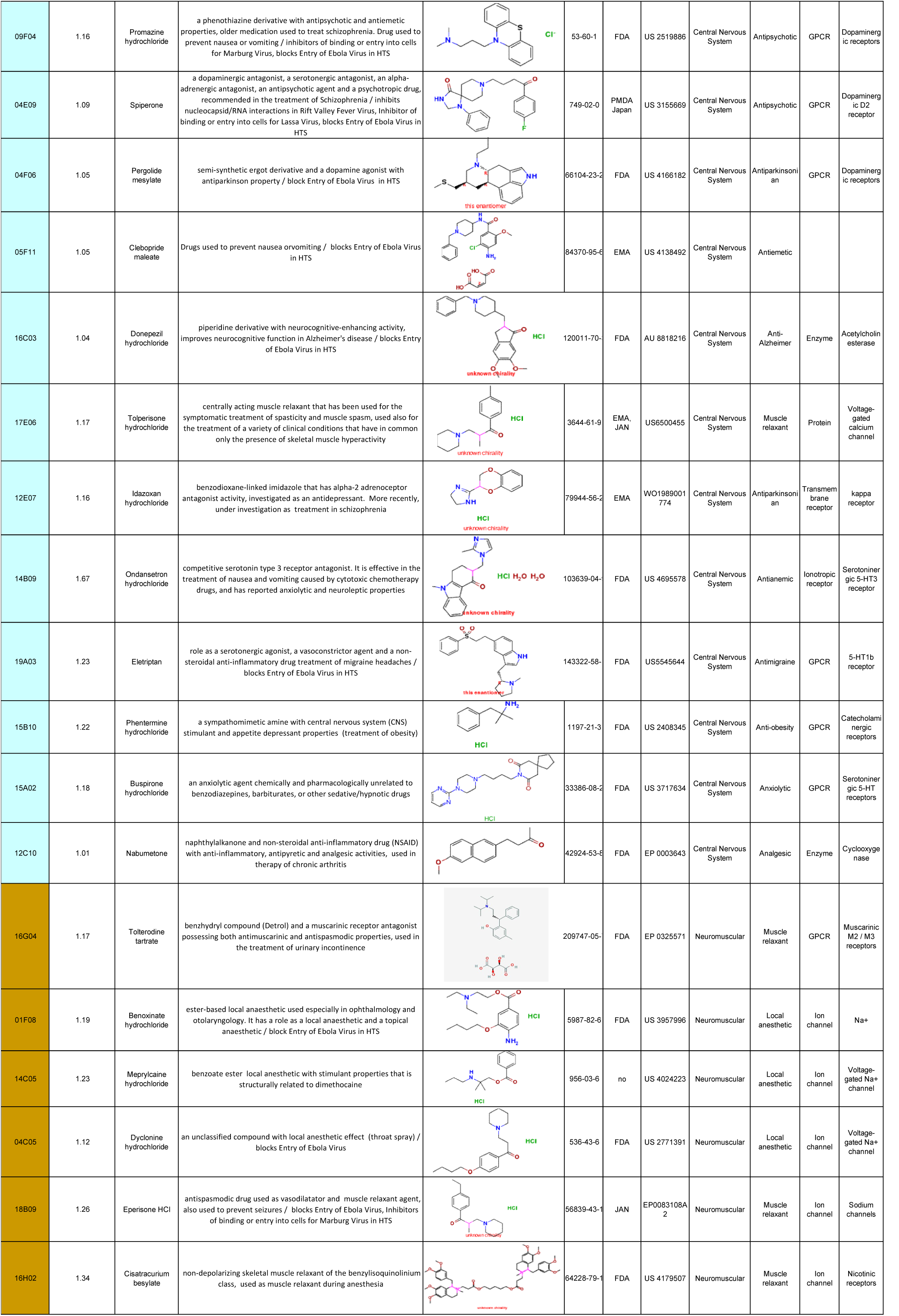
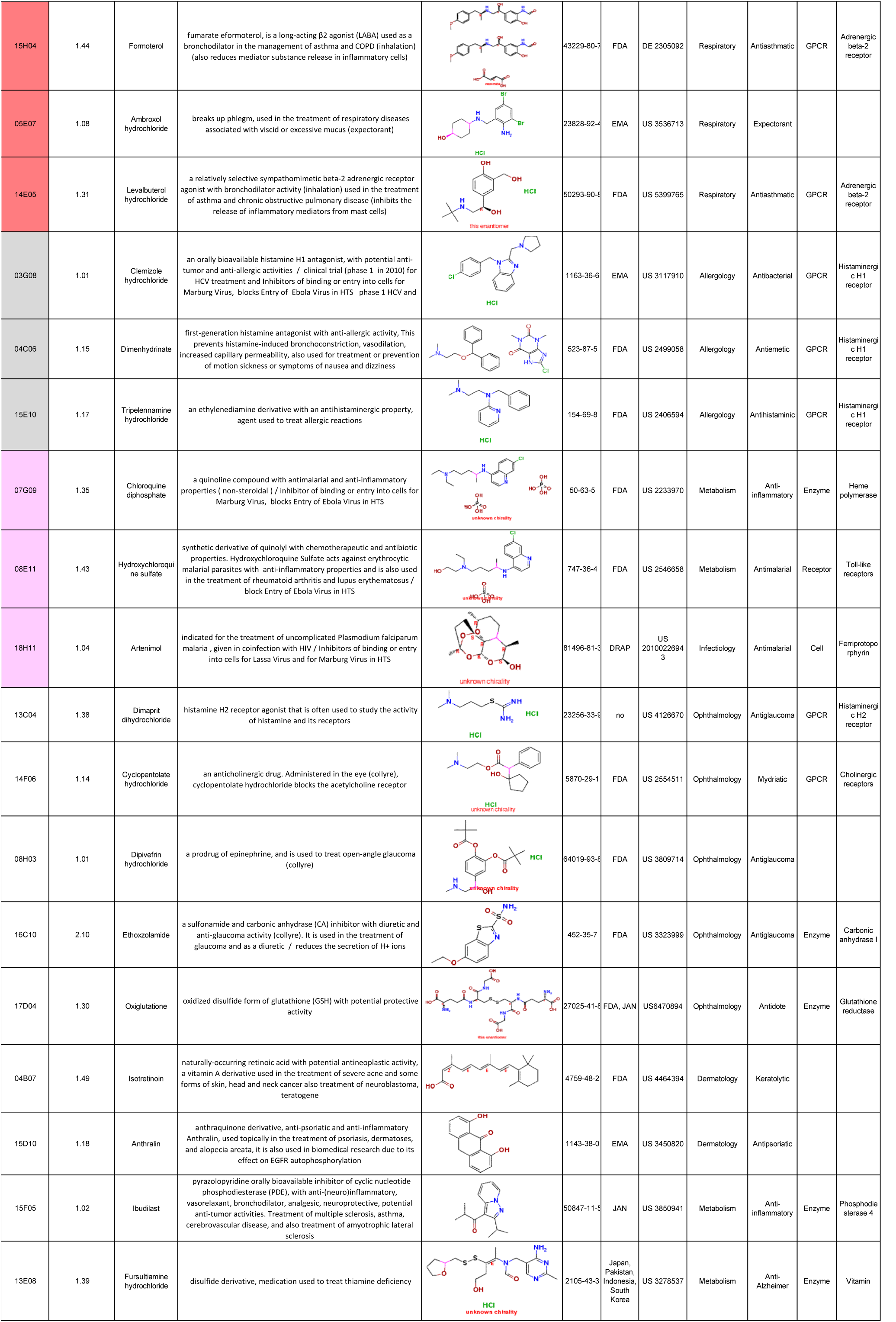
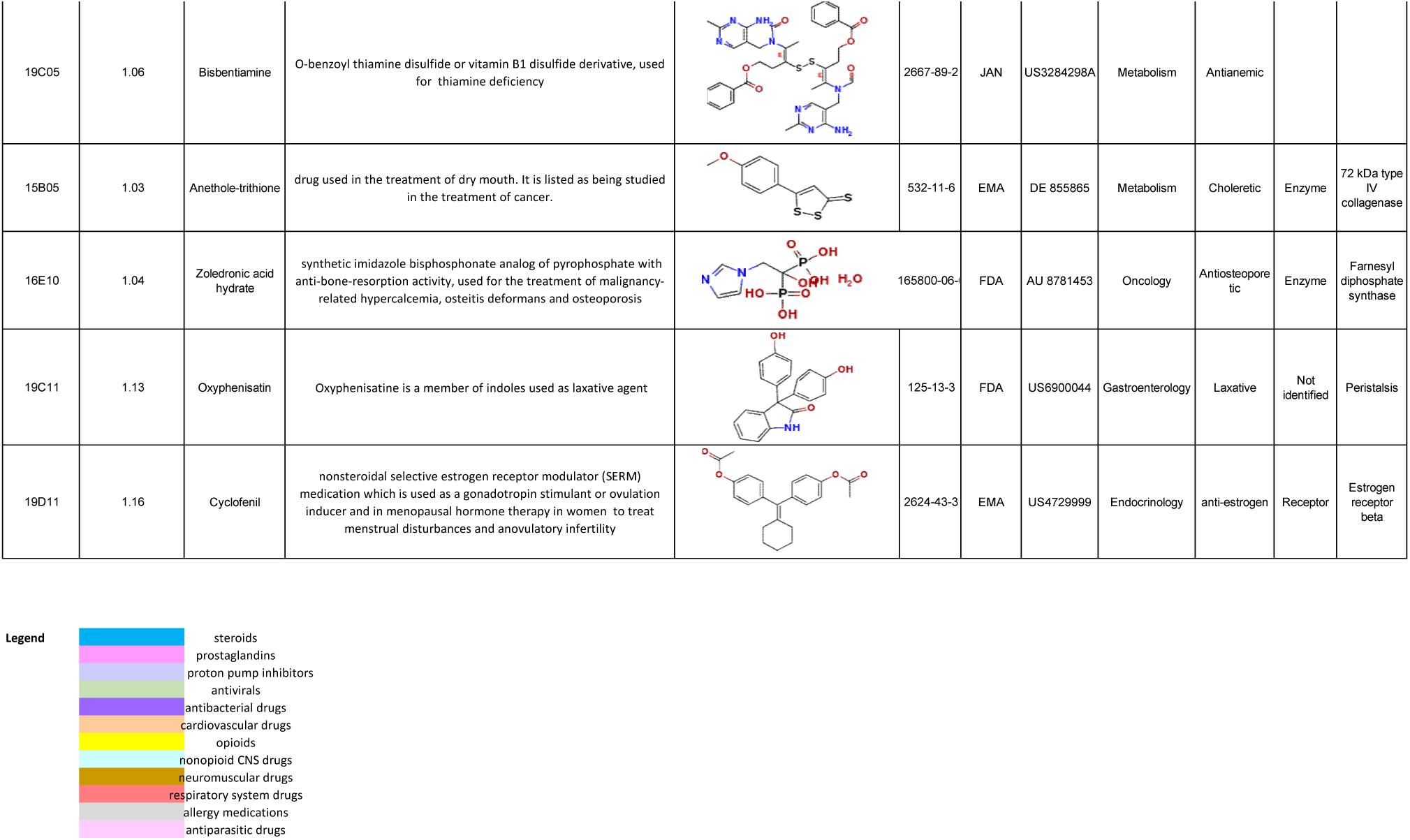
Inhibition index and detailed description of the 90 hit compounds. The compounds are organized in functional and/or structural class.

Interestingly, based on the SARS-Cov-2 infection cycle, one can infer that some of the identified molecules may inhibit specific steps of the virus replication cycle. This is illustrated for example by Candesartan, Olmesartan and Ambrisentan which interfer with angiotensin pathways, that play a key role in virus entry as the SARS-CoV2 Spike protein is known to bind to the cellular Angiotensin Converting Enzyme 2 receptor (ACE2) (Kuster et al., 2020; Lan et al., 2020). We also noted that 4 compounds (Omeprazole, Vonoprazan, Chloroquine diphosphate and Hydroxychloroquine sulfate) have been demonstrated to increase the pH of endosomial/golgian pathway either by inhibiting ATPase proton pomp, or by buffering the pH. We can thus expect that such endosomial pH modification would limit the processing of the Spike protein by endosomal proteases and in turn bloke the virus entry mediated by membrane fusion process. Finally, we also identified Darunavir, a HIV protease inhibitor might interfere with viral polyprotein processing during the replication cycle. This analysis identified at least three possible steps of the viral infection that can be targeted by approved drugs.

We next aimed at determining the EC50 of 16 drugs selected from the different groups. Whereas EC50 could be determined for 10 of them (Table 1, Fig 2), low to no antiviral effect (EC50 ≥ 40µM) was observed for Darunavir, Levalbuterol, Olmesartan, Ambrisentan and Ranolazine. As the compounds are selected from a screening based on the inhibition of the SARS-CoV-2-mediated CPE, we cannot exclude that these compounds have no antiviral effect but rather protect the cells from CPE. In addition Artenimol was rejected as it showed toxicity at 20µM concentration, in the same range of the antiviral effect. In conclusion, the screening of the approved compounds identified a set of molecules showing inhibition on SARS-CoV-2 *in vitro* replication. Some of these experimentally selected candidates, with EC50 at the 2-20 µM range may provide information to guide the choice for downstream experiments and validation, or initiate medicinal chemistry projects to find more potent derivatives.

**Figure 2:**
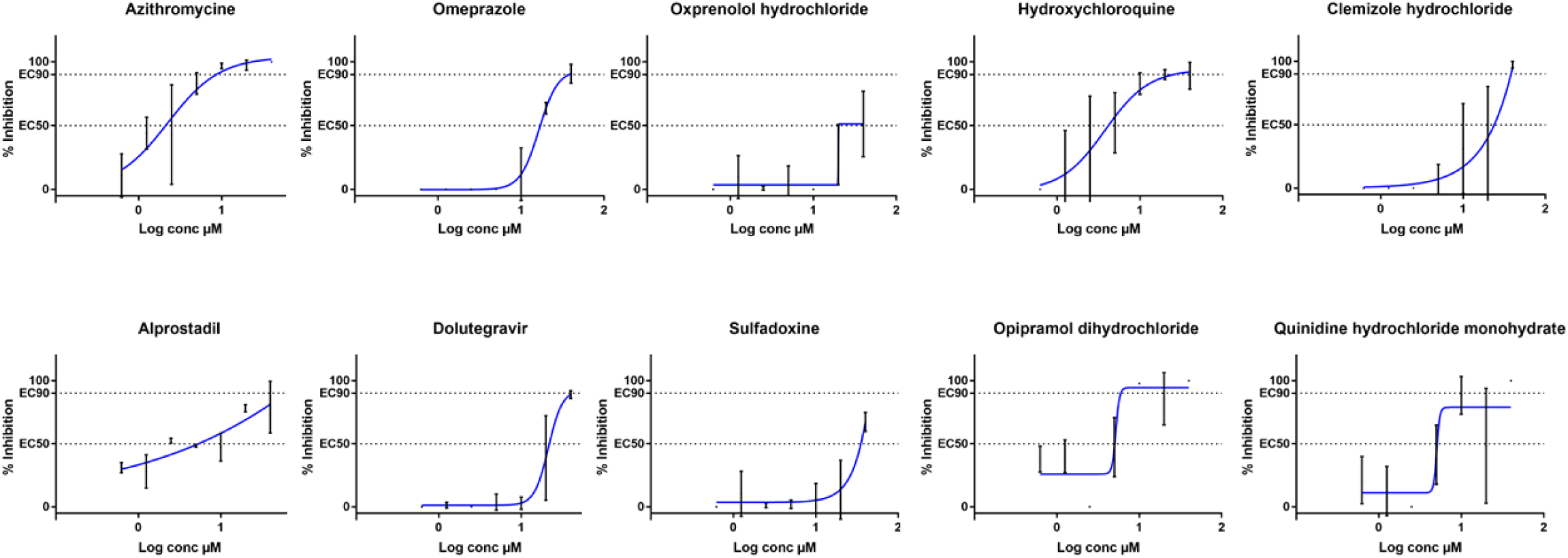
Dose response curves of selected compounds from the hits and control compounds. EC50: 50% inhibition, EC90: 90% inhibition. Compounds concentrations are in presented in log scale for logarithmic interpolation.

## Supporting information

Supplemental table 1 and raw data

## Acknowledgments

This work was supported by Inserm through the REACTing (REsearch and ACTion targeting emerging infectious diseases) initiative and by the European Virus Archive Global (EVA GLOBAL) funded by the European Union’s Horizon 2020 research and innovation programme under grant agreement No 871029. The antiviral screening platform is affiliated to the “Très Grande Infrastructure de Recherche” ChemBioFrance network. We thank Dr Marie-Louise Jung from Prestwick-Domain Therapeutics for data mining and allowing the disclosure of the information on the Prestwick Chemicals Library®, as well as Pr Drosten and Pr Drexler for providing the SARS-CoV-2 through EVA GLOBAL. We would also thank Camille Placidi-Italia for excellent technical support.

